# Automated prediction of site and sequence of protein modification with ATRP initiators

**DOI:** 10.1101/2022.07.23.501081

**Authors:** Arth Patel, Paige N. Smith, Alan J. Russell, Sheiliza Carmali

## Abstract

One of the most straightforward and commonly used chemical modifications of proteins is to react surface amino groups (lysine residues) with activated esters. This chemistry has been used to generate protein-polymer conjugates, many of which are now approved therapeutics. Similar conjugates have also been generated by reacting activated ester atom transfer polymerization initiators with lysine residues to create biomacromolecular initiators for polymerization reactions. The reaction between activated esters and lysine amino groups is rapid and has been consistently described in almost every publication on the topic as a “random reaction”. A random reaction implies that every accessible lysine amino group on a protein molecule is equally reactive, and as a result, that the reaction is indiscriminate. Nonetheless, the literature contradicts itself by also suggesting that some lysine amino groups are more reactive than others (as a function of p*K*_a_, surface accessibility, temperature, and local environment). If the latter assumption is correct, then the outcome of these reactions cannot be random at all, and we should be able to predict the outcome from the structure of the protein. Predicting the non-random outcome of a reaction between surface lysines and reactive esters could transform the speed at which active bioconjugates can be developed and engineered. Herein, we describe a robust integrated tool that predicts the activated ester reactivity of every lysine in a protein, thereby allowing us to calculate the non-random sequence of reaction as a function of reaction conditions. Specifically, we have predicted the intrinsic reactivity of each lysine in multiple proteins with a bromine-functionalised N-hydroxysuccinimide initiator molecule. We have also shown that the model applied to PEGylation. The rules-based analysis has been coupled together in a single Python program that can bypass tedious trial and error experiments usually needed in protein-polymer conjugate design and synthesis.

## Introduction

The addition of polymers to protein surfaces enables researchers to tune the properties of a protein, such as stability, solubility, and activity [1, 2]. The capacity to do “polymer-based protein engineering” has helped address some of the grand challenges in the pharmaceutical industry. Several PEGylated (poly(ethylene glycol)-modified) proteins have found applications in the drug market. For example, Enzon’s Adagen is a Food and Drug Administration approved PEGylated-protein conjugate that is used in the treatment of severe combined immunodeficiency disease [3]. Currently, the only polymer modified proteins that are FDA approved are PEGylated proteins, but their overall success is driving innovation towards better alternatives for polymer-based protein engineering such as atom transfer radical polymerization (ATRP).

ATRP is a tunable approach to growing polymers from the surface of proteins [4-6]. In this approach, amino groups on the protein surface can be covalently modified with small initiator molecules, which then serve as initiation sites for polymer growth, allowing the synthesis of an array of conjugates with different polymer compositions and architectures. Compared to standard PEGylation, ATRP provides greater control over the degree of modification and allows for higher grafting densities. Depp et al. demonstrated that ATRP-enhanced chymotrypsin had increased stability and activity retention relative to PEGylated chymotrypsin [7]. In a more recent study, the negative impact of PEGylation on activity and stability was discussed, along with alternatives to overcome them [8]. Over time, our breadth of knowledge for synthesizing and analysing the outcome of reacting activated esters with the surface of proteins has advanced considerably. Nonetheless, predicting the sequence and outcomes of these reactions has remained elusive, even for the coupling of ATRP-initiators to the surface of a protein. In general, scientists perform costly and laborious stochastic experiments to generate conjugates with the desired grafting densities and polymer-modification sites. The ability to predict how proteins are modified would greatly improve the efficiency of polymer-based protein engineering.

Over the past decade, our laboratories have studied an array of protein-polymer conjugates synthesized using ATRP [9-13]. More recently, we have shown that the tertiary structure of a protein can be used to assess the relative reactivity of each lysine in a protein. We developed a decision tree based upon experimental data, which we then used to predict the relative reactivity of amino groups (N-terminus and lysine residues) in a protein and an ATRP initiator [14]. This prediction incorporated calculated surface accessibility, p*K*_a_, local charge, and the environment of the targeted amino group. Unfortunately, the required parameters were calculated with different software packages (such as Chimera [15] and Discovery Studio [16]) in a laborious and step-wise process. Once all the calculations had been performed, we then manually implemented the decision tree to “bucket” each amino group into non-reactive, slow-reacting, or fast-reacting categories.

We have now built a Python program that uses a PDB file to predict the outcome of protein-labelling reactions in a single step, thereby automating the entire process. The program can be used to determine which lysine residues are likely to be labelled first in any protein with a known structure. We can also predict which amino groups will never be labelled. We have named this program “PRELYM” (Prediction of Lysine Modification”). Herein, we compare the output of the manual and automated predictions for lysozyme, chymotrypsin, glucose oxidase and avidin.

## Materials and Methods

### Prediction of exposed surface area

Our PRELYM Python program adapted the Lee-Richards algorithm to calculate the exposed surface area (ESA) of lysine residues and N-termini for a given protein [17]. This approach calculates the ESA by probing the surface of the molecule with a sphere having a radius equivalent to the solvent radius. For this calculation the program integrated three Python libraries; BioPandas, BioPython, and FreeSASA. The tertiary structure of the desired protein was first downloaded from the RCSB Protein Data Bank in a PDB format and then uploaded to BioPandas where the PDB file was read to find atoms associated with their respective residues [18]. Subsequently, BioPython created the BioPDB structure required to calculate the ESA [19]. This BioPDB structure along with the user input of the probe radius of the chosen initiator molecule was then passed to FreeSASA which calculated the ESA for each protein atom and reported the value in Å^2^ [20]. The ESA for each lysine residue was then calculated by adding the ESA of each atom present in that residue.

### Prediction of p*K*_a_ for protein amino groups

We designed our program to employ PROPKA to predict the p*K*_a_ for each amino group in lysine residues and the N-termini of a protein [21]. PRELYM implemented the PROPKA 3 fast heuristic p*K*_a_ predictor to calculate p*K*_a_ from protein conformations [22]. The program first used the MDAnalysis library, which takes the PDB file as an input to create an MDAnalysis Class Universe that contained all the necessary information to calculate p*K*_a_ (such as residue, atom, and atom coordinates that describe the protein structure) [23]. The Universe was then used as an input by the Python Library, PropkaTraj, to calculate the p*K*_a_ values for protein residues using the PROPKA method [24].

### Prediction of secondary structure

We used the Define Secondary Structure of Proteins (DSSP) algorithm to identify the secondary structures in which specific residues were located. The DSSP algorithm pinpointed the secondary structures through atomic coordinate analysis [25]. DSSP categorized secondary structures into 8 subcategories, from which our program then classified them into 3 categories: helix, strand, and coil. The program used a Python library, structural systems biology (ssbio) [26], that included wrappers to the BioPython DSSP Module [19] which were used to execute and obtain the DSSP results.

### Prediction of involvement of amino group in H-Donor interactions

To identify the amino groups acting as H-bond donors, we first identified the likely hydrogen bonds via the Baker-Hubbard method [27] based on cut-offs for the donor-H-acceptor angle and distances using the MDAnalysis Library [23] which had a pre-built code for hydrogen bond identification using the Baker-Hubbard algorithm. The resulting output contained hydrogen bond details in the form of donor-acceptor, from which the donor residues were extracted from a PDB file containing the hydrogens. Missing hydrogens in the PDB file were added by using MolProbity [28].

### Prediction of areas of limited charge

In the program, local charge calculations were performed using Coulomb’s Law to calculate the electrostatic energy of individual amino acids. We used a PQR file as an input which had been generated by using the PDB2PQR suite included in the APBS software [29, 30]. The PDB2PQR software applied a force field to assign charges to atoms and returned the results in the form of a PQR file. Our program then used this PQR file to calculate the electrostatic energy on each atom. The formula for this calculation was derived from Zhou et al [31]. The electrostatic energy value for the terminal zeta (NZ) position of the lysine group was considered, with a final value greater than 100 kcal/mol being considered an area of lower positive charge.

### Data processing

The parameters discussed above were calculated individually and then stored in separate Pandas DataFrames [32]. Our program then further sorted the data in a single DataFrame. First, the program identified the N-termini and lysine groups of the protein from the PDB file using BioPandas [18], and then looped over the parameter data, finding the values of each parameter for a specific residue, and adding those values to the respective parameter columns. Once the final DataFrame contained every parameter for all lysine and N-termini, the decision tree from Carmali et al. [14] was applied to the compiled data. Using the decision tree, the program predicted the reactivity of the isolated amino groups with an ATRP initiator molecule and subsequently classified each amino group as non-reacting, slow-reacting, or fast-reacting (Fig 1). It is important to note that the calculations performed are not identical to those of Carmali et al [14], and therefore we expected to observe differences in the output.

**Fig 1.**
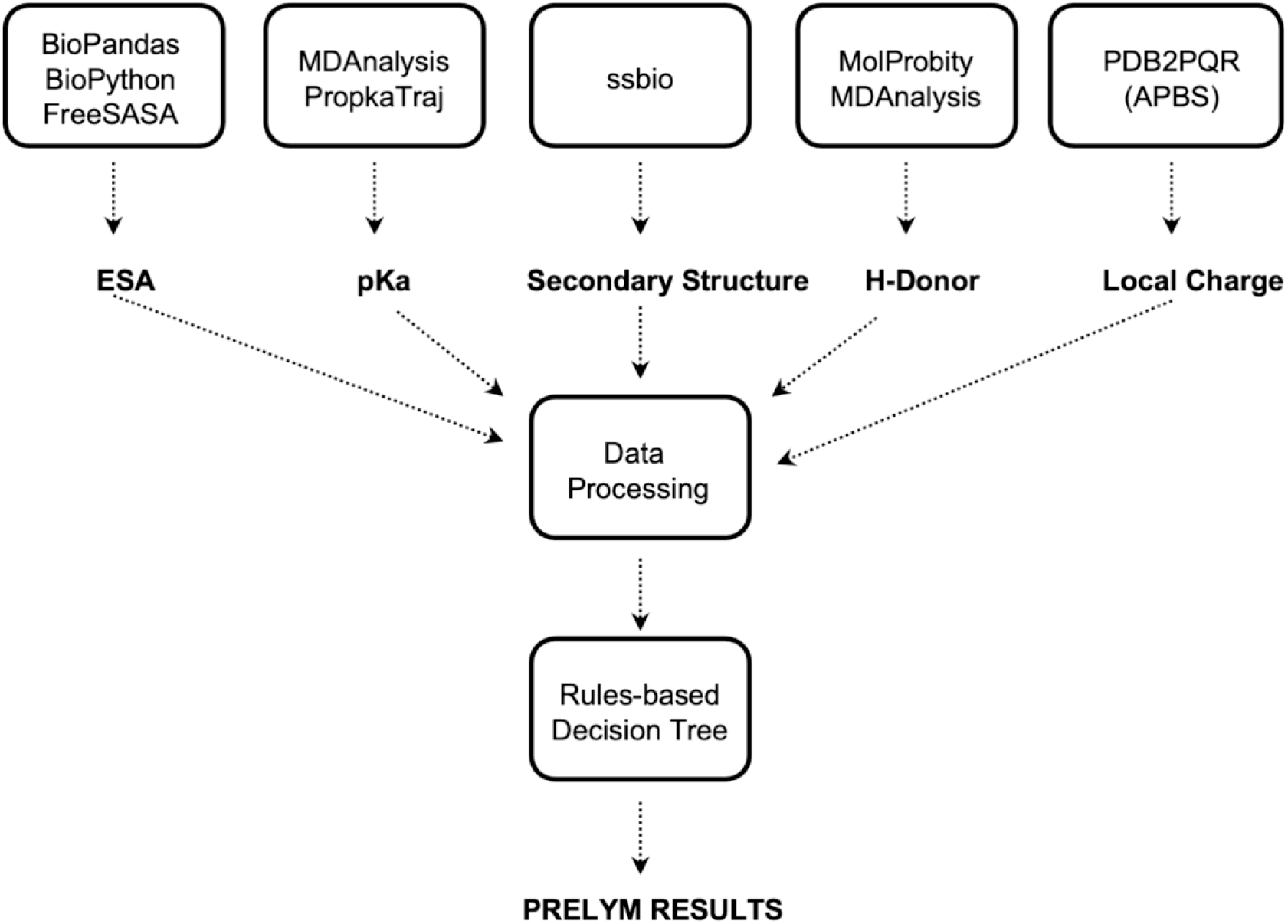
Flowchart for parameter calculation and processing using different libraries and software.

## Results and Discussion

### PRELYM: implementation and validation

To assess the accuracy of PRELYM, we first used the program to predict the reactivity of amino groups in hen egg-white lysozyme (PDB ID: 1LYZ) and alpha chymotrypsin (PDB ID: 4CHA) with a bromine-functionalised N-hydroxysuccinimide ATRP initiator. Prior to executing PRELYM, each PDB file was processed accordingly to remove water molecules, add missing atoms, and generate a PQR input file for electrostatic calculations (S1 File). The probe radius was set to 4.2 Å to match the theoretical diameter of the initiator molecule and the virtual pH set to 8.0 to mirror experimental conditions implemented in our previous work [1]. We ran the program on a Windows operating system, Intel Core i9 2.3 GHz with 16 GB RAM.

### Lysozyme

In a previous study, we determined that five out of seven possible amino sites present on the surface of lysozyme were modified with the ATRP initiator [6]. Accordingly, PRELYM accurately classified the amino reactivities of lysozyme as defined by our rules-based decision tree [14]. The program was rapid and efficient, taking only about 3 seconds to perform the calculations.

Fig 2 depicts the structure of lysozyme with the reactivities of each amino group. The complete data set obtained from PRELYM along with the predicted and experimental reactivities are summarized in S1 Table. Residues K13, K33 and K97 have an exposed surface area greater than 50 Å^2^, which according to our rules-based decision tree determines these residues as reactive. Since these residues are part of the less accessible helical fold in lysozyme [33], PRELYM accurately assigned them as slow-reacting groups. In contrast, K96 has an underexposed surface area of 26.37 Å^2^ and was classified as a non-reacting amino group. Interestingly, in the PRELYM analysis, the N-terminus lysine (K1) was found to have overexposed surface area of 131.44 Å^2^ and both α− and ε− amino groups were predicted to be fast-reacting. However, our previous site modification studies were only able to identify a single modification at K1.

**Fig 2.**
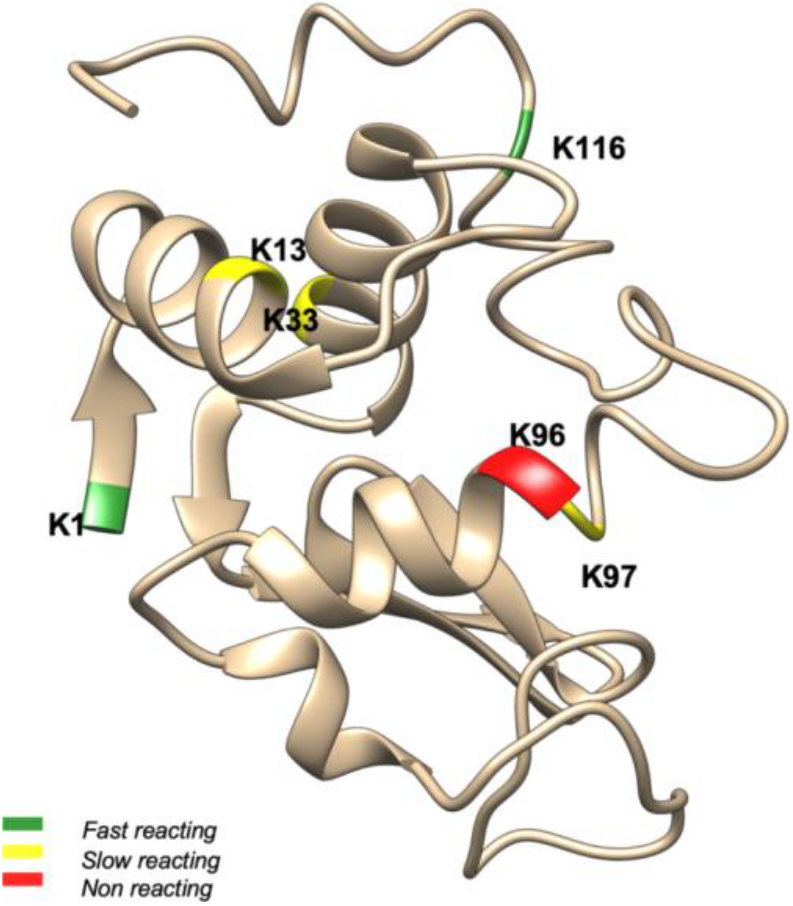
Predicted reactivities of lysozyme amino groups with ATRP initiator. The structure of lysozyme (PDB ID: 1LYZ) is presented using Chimera [15]. Lysine residues and N-termini are highlighted in green, yellow, and red to correspond to fast reacting, slow reacting, and non-reacting amino groups respectively.

Noteworthy, we also examined how PRELYM performed for bioconjugates prepared by reversible addition-fragmentation chain transfer (RAFT) polymerisation. As with ATRP, surface-initiated RAFT requires first the attachment of small molecule compounds, called chain transfer agents (CTA), which serve as starting points for polymer growth [34]. Here, we selected an *N*-hydroxysuccinimide ester CTA containing a diethylene glycol spacer [35] and an oligomeric CTA composed of 5 acrylamide units [36]. Both CTAs have been used to generate lysozyme-polymer conjugates with preferential modifications at K1, K33 and K97 [35, 36], matching our amine-ATRP initiator results (S1 Table). Like with the ATRP initiator reaction, these RAFT CTAs were coupled to lysozyme using N-hydroxysuccinimide chemistry, and as such, we did not expect significant differences. To better resemble lysozyme-CTA interactions, we also executed PRELYM with the probe radius reflecting the approximate size of each CTA. For this, we first estimated the theoretical diameter of each CTA molecule, by measuring end-to-end distances (S4 Fig). Predicted amine reactivities using the new CTA probe radii (S2-3 Tables) were overall consistent with our initial PRELYM results using the 4.2 Å probe (S1 Table). Discrepancies in relative reactivities were observed but these can be attributed to variations in experimental reaction conditions. For example, lysozyme reactions with oligomeric CTA included *in situ* active ester formation which could influence modification [36]. Nevertheless, this highlights how PRELYM can be broadly applied to predict protein-amine reactivity with small-molecule active esters.

### Chymotrypsin

We next confirmed the accuracy of PRELYM using the crystal structure of bovine dimeric α – chymotrypsin (PDB ID: 4CHA). After generation of the required input files, PRELYM results were obtained after approximately 13 seconds (S5 Table). This slight increase in computational time was driven by the relative sizes of chymotrypsin and lysozyme. Homo-dimerization of chymotrypsin is pH-dependent and reversible [37]. In the laboratory, ATRP initiator modifications are typically conducted under neutral to slightly alkaline pH conditions, where chymotrypsin is monomeric. We hypothesized that the amino-ATRP initiator interactions, especially those at the dimer interface, would change with the protein’s quaternary structure. To assess these variations, we modified the PDB file to depict chymotrypsin as a monomer prior to executing PRELYM (S6 Table).

Unsurprisingly, residues K36, K90, A149 and K175 present at the dimer interface showed increased accessibility in monomeric chymotrypsin (Fig 3 and S5-S6 Tables). For K90, the increased exposed surface area from 86.98 Å^2^ to 117.85 Å^2^ was accompanied by an increased pKa from 10.12 to 10.33. Given that our rules-based decision tree has a branching point classification for amino groups with pKa ≤ 10.3, PRELYM categorizes K90 as fast- or slow-reacting when present in dimeric or monomeric chymotrypsin, respectively. From our site modification studies at pH 8.0, when chymotrypsin exists as a monomer, K90 was found to be slow-reacting matching PRELYM’s results. Residue A149, the N-terminal group (chain C in sub-unit 1), in dimeric chymotrypsin is extensively buried and involved in electrostatic and hydrophobic interactions with G59, D64 and F41 (chain F in sub-unit 2). As a result, PRELYM classifies the α-amino group in A149 as non-reacting. However, in the monomer form, A149 is over-exposed (ESA 261.37 Å^2^) and predicted to be a fast-reacting amino group. These discrepancies in amino-ATRP interactions highlight the importance of understanding the quaternary structure of the protein in solution for an accurate PRELYM analysis.

**Fig 3.**
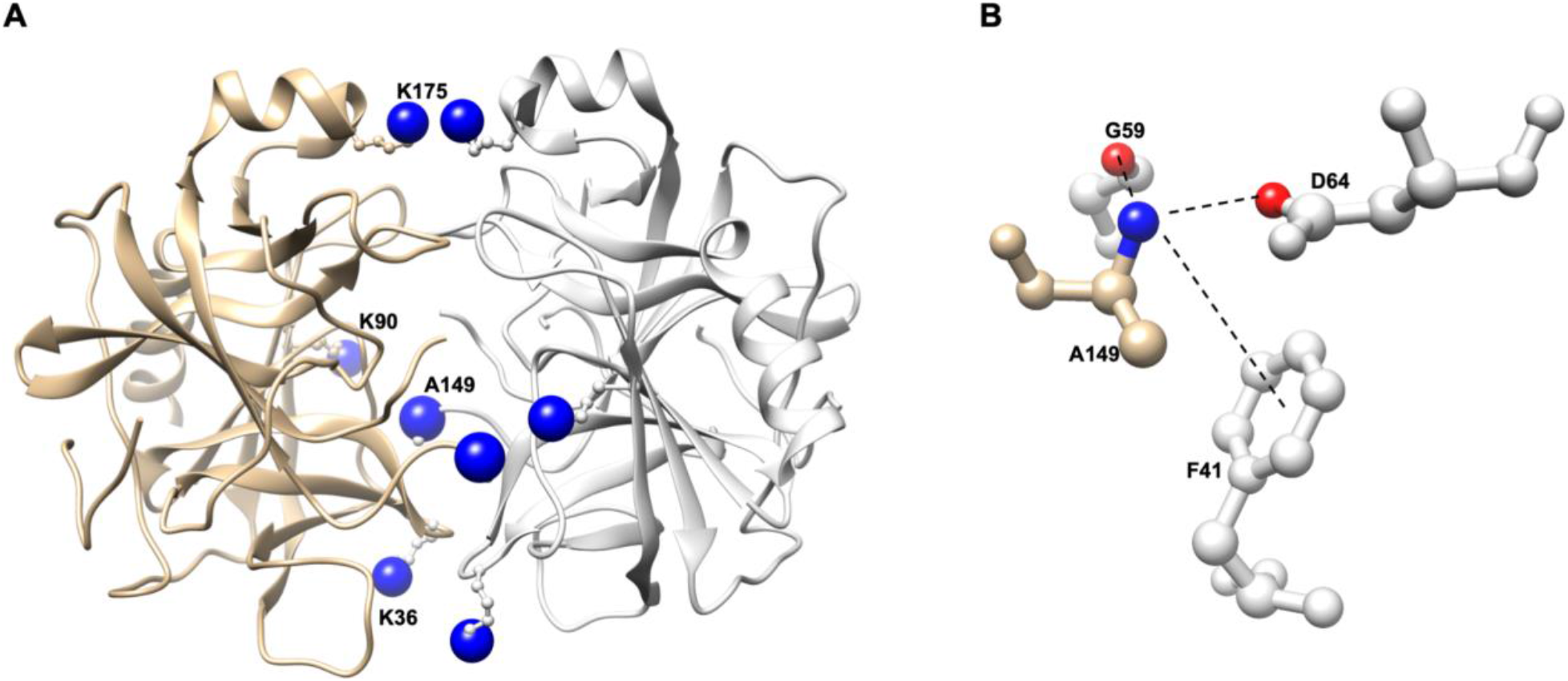
Structure of a-chymotrypsin dimer from *Bos taurus* (PDB ID: 4CHA) **A** depicting residues K36, K90, A149 and K175 at the dimer interface, with α – and ε – amino groups shown as blue spheres and each monomer sub-unit coloured tan or light grey; **B** electrostatic and hydrophobic interaction between α – amino group from A149 with G59, D64 and F41.

For the monomer of chymotrypsin, apart from residues K79 and K82, results from our program were the same as we had determined experimentally [14]. Differences in local charge values calculated for these residues caused this discrepancy.

### PRELYM for ATRP modified proteins

To further examine the capability of PRELYM, we sought to examine how PRELYM performed when predicting modification using other proteins than those previously used to generate our decision tree. For this we chose two proteins, Glucose Oxidase (GOx) and Avidin, that have previously been used in ATRP modification studies in our lab.

### Glucose Oxidase

Functionally active GOx is a homo-dimeric, -trimeric or -tetrameric protein [38]. Each GOx monomer sub-unit has 16 primary amine residues, including the N-terminus. Our previous studies have shown that upon reaction with bromine functionalised ATRP initiator, on average, 25 initiator molecules are immobilised onto the dimeric surface [39]. PRELYM analysis of GOx monomer (PDB ID: 1GAL) showed that 12 lysine residues plus the N-terminus are available to react with the ATRP initiator molecule (S8 Table). Input files for PRELYM were based on the GOx monomer structure since this was the only resolved protein structure of glucose oxidase in the Protein Databank at the time of this study. Considering GOx as a dimer, our PRELYM analysis suggests that 26 amine residues will react with the ATRP initiator, which was close to our experimental findings. The discrepancy between experimental and predicted results can be attributed to the inadequate quaternary structure representation. It is possible that one of the lysine residues may be present at the dimeric interface and therefore less reactive due to steric hindrance. As with our previous discussion with chymotrypsin, these results further highlight the importance of using a starting protein structure that corresponds to the experimentally used protein conformation to adequately predict with confidence.

### Avidin

Avidin is a homo-tetrameric protein, with each monomer sub-unit containing 9 lysine residues and 1 N-terminus [40]. Reaction of avidin with bromine functionalised ATRP initiator has been found to yield a macro-initiator complex with 8 initiator molecules immobilised per avidin monomer [41]. PRELYM analysis of avidin in the dimeric conformation (S9 Table) indicated that the N-terminus and 7 lysine residues (K3, K45, K71, K90, K94, K111 and K127) would be reactive towards the ATRP initiator, which are in alignment with experimental studies. Moreover, tryptic digestion studies of the avidin macro-initiator complex confirmed ATRP initiator immobilisation onto lysine residues K45, K71 and K111, as predicted by PRELYM.

### PRELYM for PEGylated proteins

Having used PRELYM to predict amine-initiator reactivities in surface-initiated ATRP bioconjugates, we next wanted to expand the application of PRELYM towards PEGylated proteins. For this, we investigated lysozyme, chymotrypsin, interferon-α 2a, asparaginase II, and phenylalanine ammonia lyase which have all been modified with PEG using *N*-hydroxysuccinimide chemistry.

Lee and Park previously reported K33, K97 and K116 as preferential amine sites when reacting lysozyme with a 3.4 kDa PEG at pH 8.0 [42]. Maiser and co-workers further reported K1 and K33 as the most reactive sites, with minor modifications at K13, K97, and K116 [43-45]. These findings agreed with our PRELYM results, particularly in the overall classification of reacting and non-reacting residues (S1 Table). Fig 4 shows how PRELYM’s predictions compare to confirmed amine sites after lysozyme modification with ATRP initiators [14], diethylene glycol [35] or oligomeric RAFT CTAs [36], and PEG 3.4 [42] or 5 kDa [43, 44]. Using the default 4.2 Å probe radius, PRELYM was able to correctly classify reacting/non-reacting amine sites for this series of *N*-hydroxysuccinimide labelling reagents. When focusing on relative reactivities (*i*.*e*., fast, or slow-reacting residues), discrepancies were observed, even for PEGs of similar molecular weights. We note that amine reactivity and the degree of PEGylation are often dependent on experimental conditions (pH, temperature, time, stoichiometry and coupling chemistry) [43, 44]. This is also true for chymotrypsin, where researchers have varied experimental conditions to obtain 3 – 8 PEGylated sites [46-48]. To our knowledge, PEGylation sites in chymotrypsin have not been identified, which has restricted our analysis with this protein. However, PRELYM predicted 14 reactive sites for chymotrypsin, with 6 fast-reacting sites (S6 Table) which are in line with the above experimental reports.

**Fig 4.**
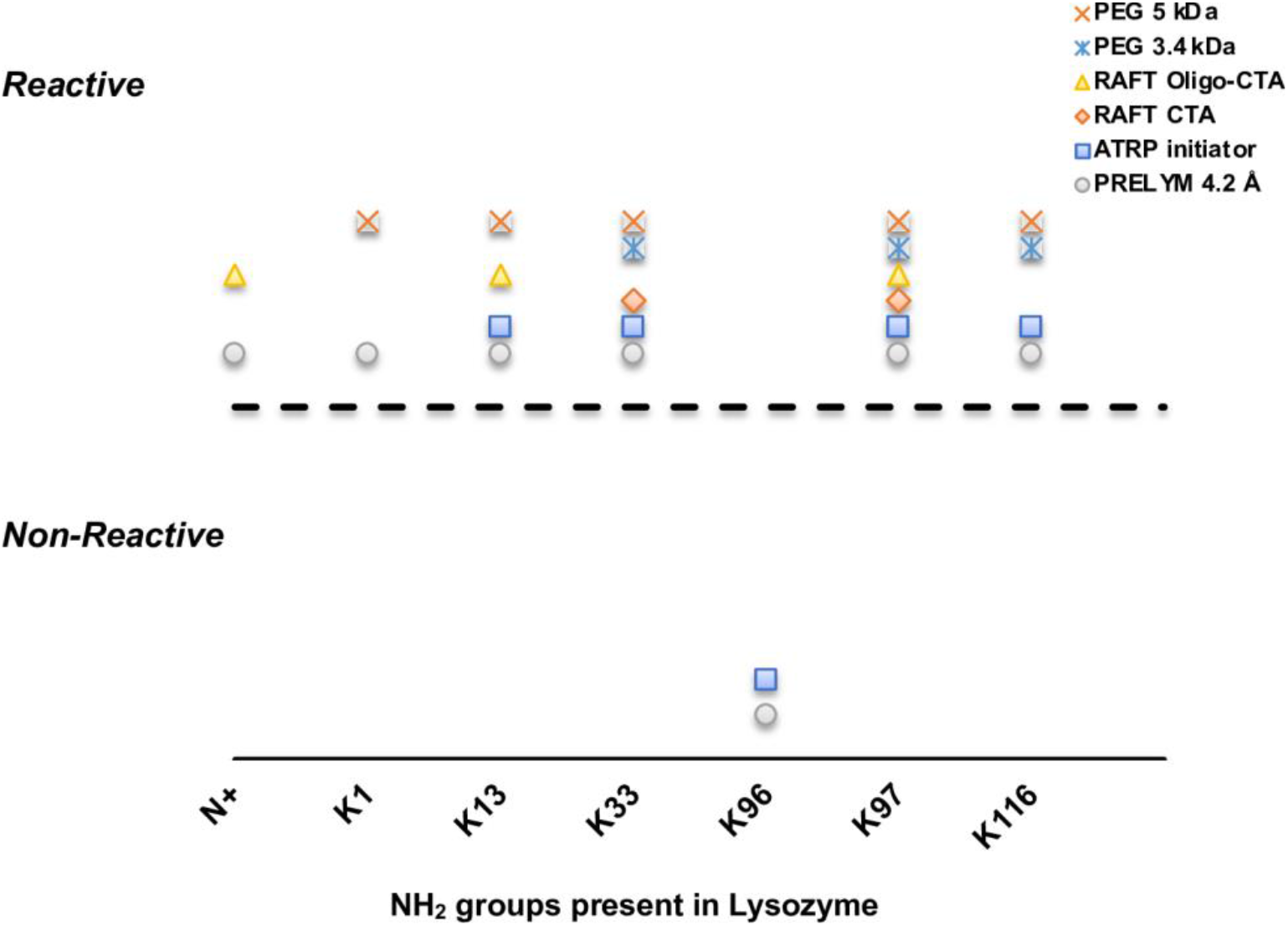
Comparison of PRELYM results using a 4.2 Å probe size with confirmed amine sites after modification with ATRP initiator [14], diethylene glycol [35] or oligomeric RAFT CTA [36], and PEG 3.4 [42] or 5 kDa [43, 44]. Overall classification of reacting and non-reacting residues in PRELYM agreed strongly with experimental findings.

Interferon-α 2a, used in the treatment of chronic hepatitis C, is formulated with a 40 kDa branched poly(ethylene) glycol chain (PEG-INF, PEGASYS) for improved half-life and efficacy. Reaction with the branched PEG has been found to afford a multi-PEGylated interferon-α 2a species with 9 positional PEG-INF isomers (K31, K49, K70, K83, K112, K121, K131, K134 and K164) [49]. Interestingly, executing PRELYM with the probe radius set to 4.2 Å matched reasonably well with experimental studies (S10 Table), with 9 lysine residues predicted to be modified. PRELYM’s prediction only diverged for the non-modified K23 and modified K112, which were predicted to be fast-reacting and non-reacting, respectively. We hypothesized that the prediction accuracy could be further improved by altering the probe size to better reflect the size of the PEG molecule used.

Thus, we repeated our PRELYM analysis using a probe radius of 39.5 Å, the equivalent to the hydrodynamic radius of PEG 40 kDa. However, this change led to decreased accuracy, with K164 predicted as a non-reacting residue (S11 Table).

Similarly, we also noted this decreased accuracy with increased probe radius with asparaginase II. Asparaginase II is a homo-tetrameric protein, with 23 primary amino groups per monomer [50]. PRELYM analysis predicted that 17 of those 23 amino groups were available for reaction, when using a 4.2 Å probe radius (S12 Table). PEG-asparaginase, when modified with amine-reactive PEG 5 kDa, leads to 69 – 82 sites modified per tetramer [51], which is in close alignment with the PRELYM’s prediction (17 sites per monomer; 68 sites per tetramer). Increasing the probe radius to 17 Å, to approximate the hydrodynamic radius of PEG 5 kDa only further decreased the number of modifications to 14 per monomer and hence, only 56 sites per tetramer (S13 Table). We hypothesized that increasing the probe size simply exacerbated the difference between which amino group is accessible or buried. For example, K107, K186 and K213 were classified as slow reacting when analysed with a 4.2 Å probe radius, with ESA < 100 Å^2^. Increasing the probe radius to 17 Å only decreased their ESA to 0 Å^2^, resulting in a non-reacting classification.

PRELYM analysis of tetrameric phenylalanine ammonia lyase (PAL) with the probe radius set to 4.2 Å was also performed. PAL is a very large protein composed of four identical sub-units used in the treatment of phenylketonuria [52]. Each monomeric sub-unit has 19 primary amine residues. PRELYM results suggested that 10 primary amines per monomer were predicted as reactive. Comparison to the experimental results with PEGylated Pal using a 20 kDa *N*-hydroxysuccinimide PEG indicated that our reactivity predictions were in good agreement (S14 Table). Moreover, PRELYM was able to correctly predict that only one of the K493/494 residue pair would be modified due to steric impediment. Discrepancies were only found with experimentally modified sites K2, K115 and K419, which were classified as non-reacting residues. We sought also to compare how an increased probe size would alter our amine reactivity predictions; however PRELYM was unable to complete surface accessibility calculations. This suggests an inherent limitation to the program, possibly based on the protein size and probe radius.

These findings indicated that increasing the probe radius did have a significant impact on the prediction and only led to changes in exposed surface area calculations that could amplify errors in PRELYM prediction. Fig 5 compares the outcome of PRELYM for each lysine residue present in interferon-α 2a when varying the probe radius. In this analysis, we selected four different probe radii, to resemble the ATRP initiator, PEG 10, 20 and 40 kDa molecules (4.2, 22.9, 30.1 and 39.5 Å, respectively). We observed that PRELYM performed with the 4.2 Å probe radius was sufficient to provide a good indication of the probability of the amine – *N* -hydroxysuccinimide reaction with proteins, even for ‘grafting to’ strategies, such as PEGylation. Discrepancies in reactivity prediction for K23 and K112 were still observed with increased probe size. Furthermore, larger probe radii led to an additional error prediction, with K164 classified incorrectly as a non-reacting residue. This can be understood by considering how exposed surface area (ESA) is defined and its dependence on probe size [53]. ESA is the molecular surface area that is created by tracing the center of a probe sphere as it rolls along the van der Waals surface. With increased probe size, ESA values also increase, due to the increased distance between the center of the probe sphere and the atoms’ center. We observed this increased ESA in lysozyme, chymotrypsin, interferon-α 2a, asparaginase II and phenylalanine ammonia lyase (S4, S7, S10-S14 Tables). However, with increased size, the probe will also have fewer spaces in the interior and crevices of the protein to fit, leading to lower ESA values for residues located more internally, but not buried (e.g., K107, K186 and K213 in phenylalanine ammonia lyase). Moreover, this large probe effect will also create a smoother molecular envelope, which can erroneously attribute residues with zero exposed surface area. This is the case for K164 in interferon-α 2a, where the increased probe radius led to the abrupt transition in classification from fast-reacting to non-reacting (Fig 5 and S10-S11 Tables).

**Fig 5.**
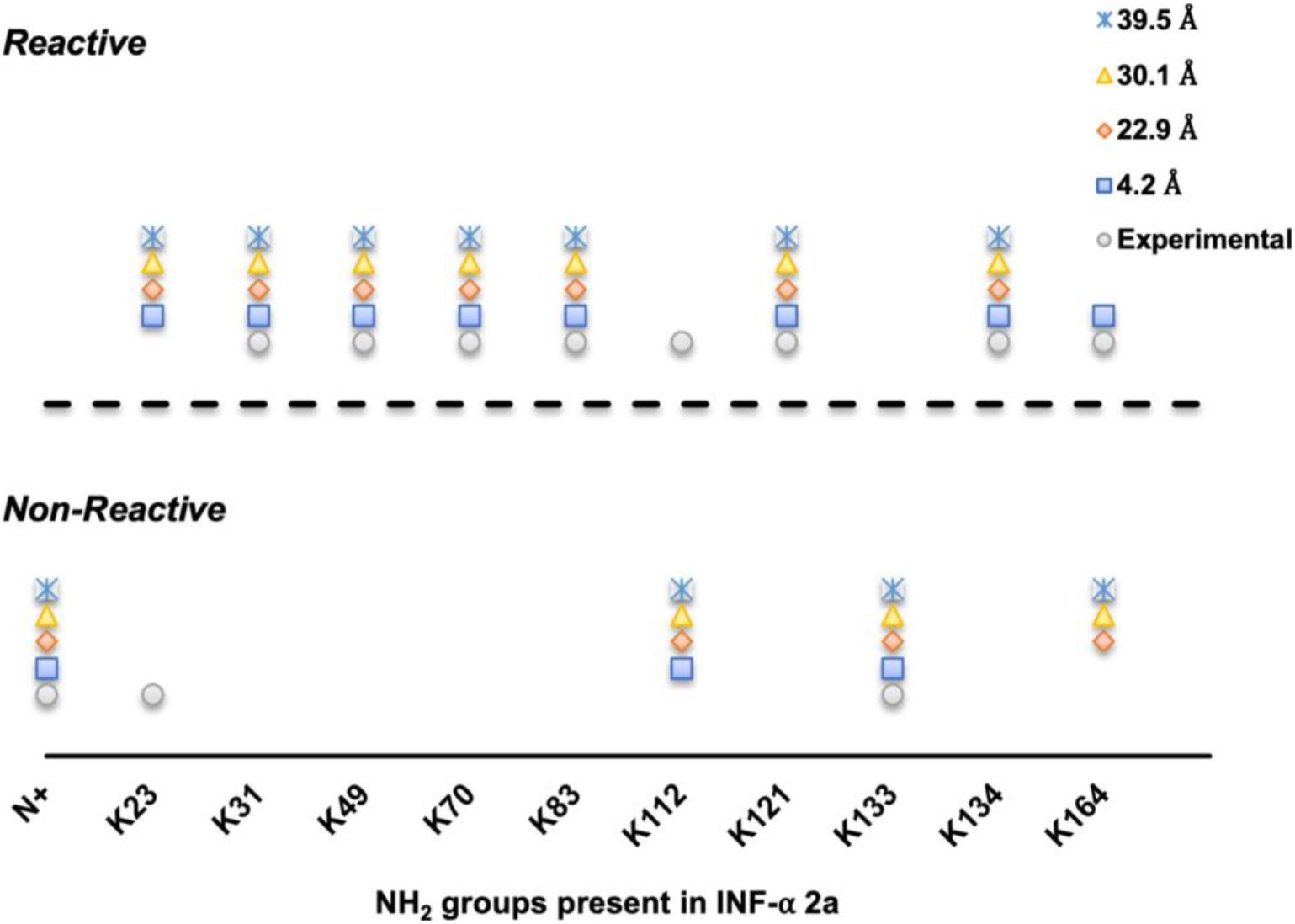
Comparison of PRELYM analysis of amine reactivities in interferon-α 2a with increasing probe radius. Increased probe radius does not improve amine reactivity prediction when considering PEGylation reactions and can lead to errors in amine residue classification.

## Conclusions

Herein, we examined the use of PRELYM to predict the reactivities of amino groups in chymotrypsin, lysozyme, glucose oxidase and avidin when reacting with an *N*-hydroxysuccinimide ATRP initiator molecule. Results obtained were found to be in good agreement with those found experimentally. PRELYM computational time was dependent on protein size and accuracy was improved when considering protein quaternary structure. We further explored the use of PRELYM in understanding protein interactions with amine-reactive PEG molecules. Increased probe size was used to resemble the hydrodynamic volume of PEG. PRELYM was also able to predict PEGylation sites in interferon-α 2a, asparaginase II and phenylalanine ammonia lyase when using a 4.2 Å probe radius. Increasing probe radii did not improve amine reactivity prediction and led to errors in PRELYM reactivity classification. Further improvements in PRELYM prediction accuracy may require the incorporation of an additional parameter to consider polymer dynamics in solution and steric effects when covalently attached to the protein. PRELYM is a unique computational tool that can be used to predict amine reactivities in bioconjugation reactions. This can provide experimental scientists with the ability to optimize reaction conditions for the synthesis of the bioconjugate they prefer, as well as improving our understanding of location-function relationships in protein-polymer conjugates.

## Supporting Information

**S1 File How to run PRELYM**.

**S1 Fig. Folder view with PRELYM input and output files**.

**S2 Fig. Creation and activation of local environment**.

**S3 Fig. PRELYM execution using Python 3.7.4 in the Anaconda Prompt**.

**S4 Fig. Chain transfer agents used for surface-initiated RAFT of lysozyme**.

**S1 Table PRELYM results for amine-ATRP initiator interactions on the surface of lysozyme**.

**S2 Table PRELYM results for amine interactions on the surface of lysozyme using a probe radius equivalent to the hydrodynamic radius of N-hydroxysuccinimide RAFT CTA (7 Å; see S4 Fig.)**.

**S3 Table PRELYM results for amine interactions on the surface of lysozyme using a probe radius equivalent to the hydrodynamic radius of oligomeric RAFT CTA (8.8 Å; see S4 Fig.).**

**S4 Table PRELYM results for amine interactions on the surface of lysozyme using a probe radius approximate to the hydrodynamic radius of PEG 5 kDa (17 Å)**.

**S5 Table PRELYM results for amine-ATRP initiator interactions on the surface of homo-dimer chymotrypsin**.

**S6 Table PRELYM results for amine-ATRP initiator interactions on the surface of monomer chymotrypsin**.

**S7 Table PRELYM results for amine interactions on the surface of monomer chymotrypsin using a probe radius approximate to the hydrodynamic radius of PEG 5 kDa (17 Å)**.

**S8 Table PRELYM results for amine-ATRP initiator interactions on the surface of monomer glucose oxidase**.

**S9 Table PRELYM results for amine-ATRP initiator interactions on the surface of dimer Avidin**.

**S10 Table PRELYM results for amine-ATRP initiator interactions on the surface of interferon-α 2a**.

**S11 Table PRELYM results for amine interactions on the surface of interferon-α 2a using a probe radius equivalent to the hydrodynamic radius of PEG 40 kDa (39.5 Å)**.

**S12 Table PRELYM results for amine-ATRP initiator interactions on the surface of asparaginase II**.

**S13 Table PRELYM results for amine interactions on the surface of asparaginase II using a probe radius approximate to the hydrodynamic radius of PEG 5 kDa (17 Å)**.

**S14 Table PRELYM results for amine-ATRP initiator interactions on the surface of tetrameric phenylalanine ammonia lyase (rAV-PAL)**.

