## Supporting Information for "Automated prediction of site and sequence of protein modification with ATRP initiators"

**Installing Anaconda as a pre-requisite in Windows OS.** To run PRELYM, we first installed Anaconda, to create a virtual environment. The 64-Bit Graphical Installer for Anaconda was downloaded from [www.anaconda.com/products/individual#windows](http://www.anaconda.com/products/individual#windows). On-screen instructions were followed and upon successful installation, the Anaconda prompt will be available in the search bar.

**Preparation of input files for PRELYM.** To execute PRELYM, three input files are necessary. As an example, the process of preparing these files will be explained using the protein Avidin. The PDB file for Avidin (PDB ID: 2AVI) was downloaded from the Protein Databank ([www.rcsb.org](http://www.rcsb.org)). Removal of water molecules and the addition of missing atoms was accomplished using Discovery Studio. The modified PDB file was then saved as a new file, named 2AVI_modified.pdb, and was the first input file.

To generate the second input file, we used the previously prepared first input file and MolProbity (<http://molprobity.biochem.duke.edu>). MolProbity is a general-purpose web server that offers quality validation for 3D structures of proteins, nucleic acids, and complexes[1]. For this study, MolProbity allowed us to add and fully optimise bond lengths for all hydrogen atoms present in the protein structure. Once this process was completed, we obtained a new PDB file with the optimised hydrogen atoms named 2AVI_modifiedFH.pdb.

For the third input file, we prepared a .pqr file using the APBS server (<http://server.possionboltzmann.org>), using the PDB2PQR job configuration tool. Here, the first input file was used, and appropriate settings for pH were selected. In our study, we used pH 8.0 to be aligned with the experimental conditions used in previous studies. Prior to initiating the process, we confirmed that the ‘create an APBS input file’ was selected. Upon completion, a list of files was generated. As the third input, the PQR file was downloaded and renamed to 2AVI.pqr.

**Executing PRELYM.** To execute PRELYM, the three input files generated previously along with the two program files (.py and .yml, freely available for download at GitHub <https://github.com/scarmali/PRELYM> ) were placed in a common folder (S1 Fig).

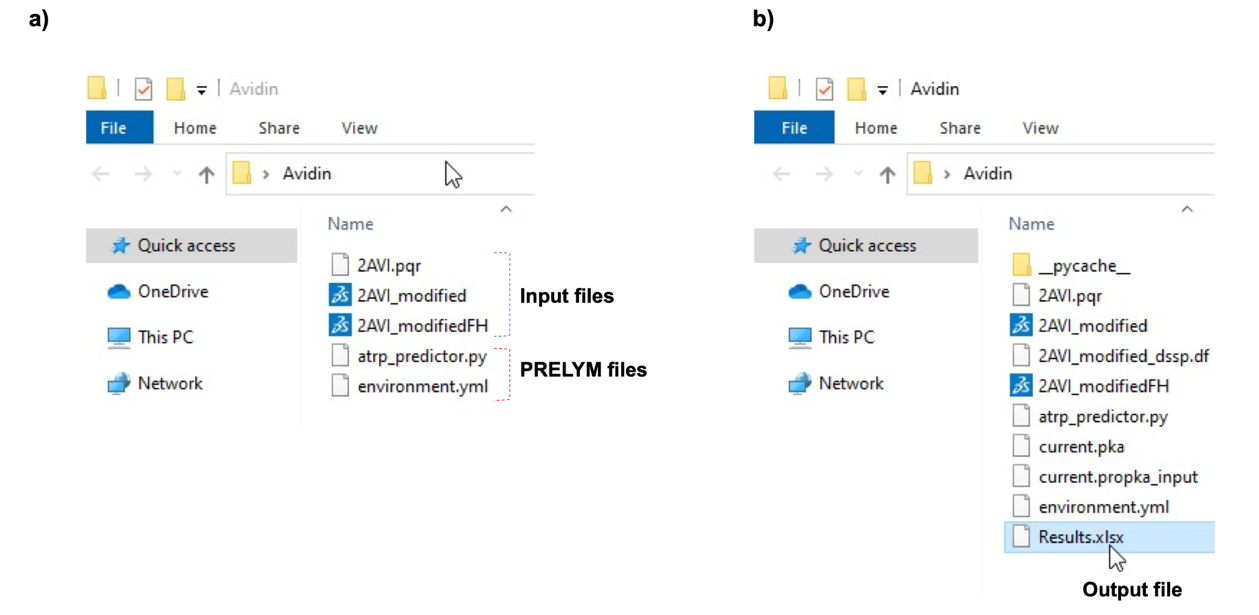

S1 Fig. Folder view with PRELYM input and output files. In the common folder a) three generated input files and the two PRELYM program files are placed for PRELYM execution; b) after running PRELYM, a ‘results’ spreadsheet is generated as the output.

In the Anaconda Prompt, the current working directory was set to the path where the common folder with the required files was located. The next step involved creating a virtual environment for the local computer and is only necessary to be done once. In the Anaconda Prompt, *conda env create -f environment.yml* was typed. When prompted, *y* was selected and upon completion, a prompt to either activate or deactivate conda appeared. Conda activation led to a change from base to ATRP environment, indicating the successful creation of the local environment (S2 Fig). Python was then used in Anaconda to execute PRELYM. In the Anaconda prompt, PRELYM was called by typing *atrp_predictor import decision_tree* in the command line. The function *decision_tree* was then provided with four arguments: the three input files generated previously, and the selected probe radius (S3 Fig). For ATRP initiator modifications, the probe radius was set to 4.2 Å. PRELYM will was then run for a period dependent on the protein size. Once completed, a ‘Results.xlsx’ output file was automatically generated in the common folder (S1 Fig).

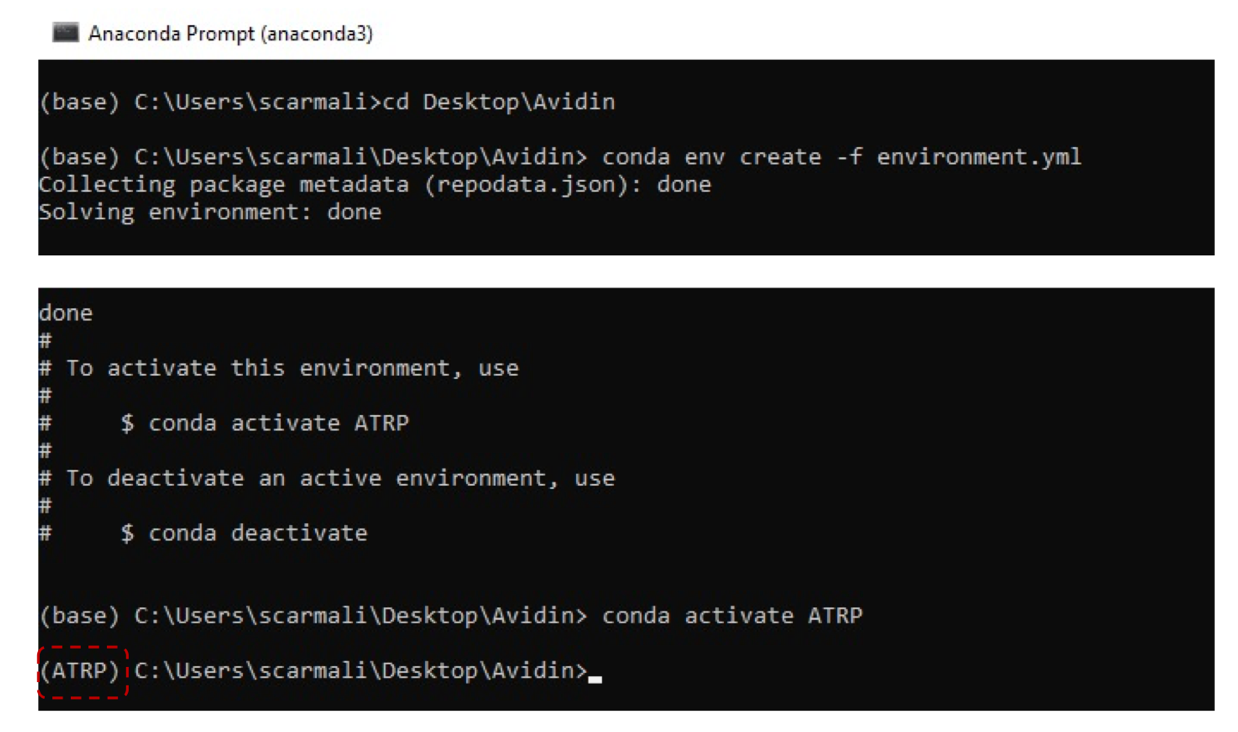

S2 Fig. Creation and activation of local environment. Anaconda Prompt showing creation and activation of local environment. Highlighted in red indicates the successful activation of the local environment from base to ATRP.

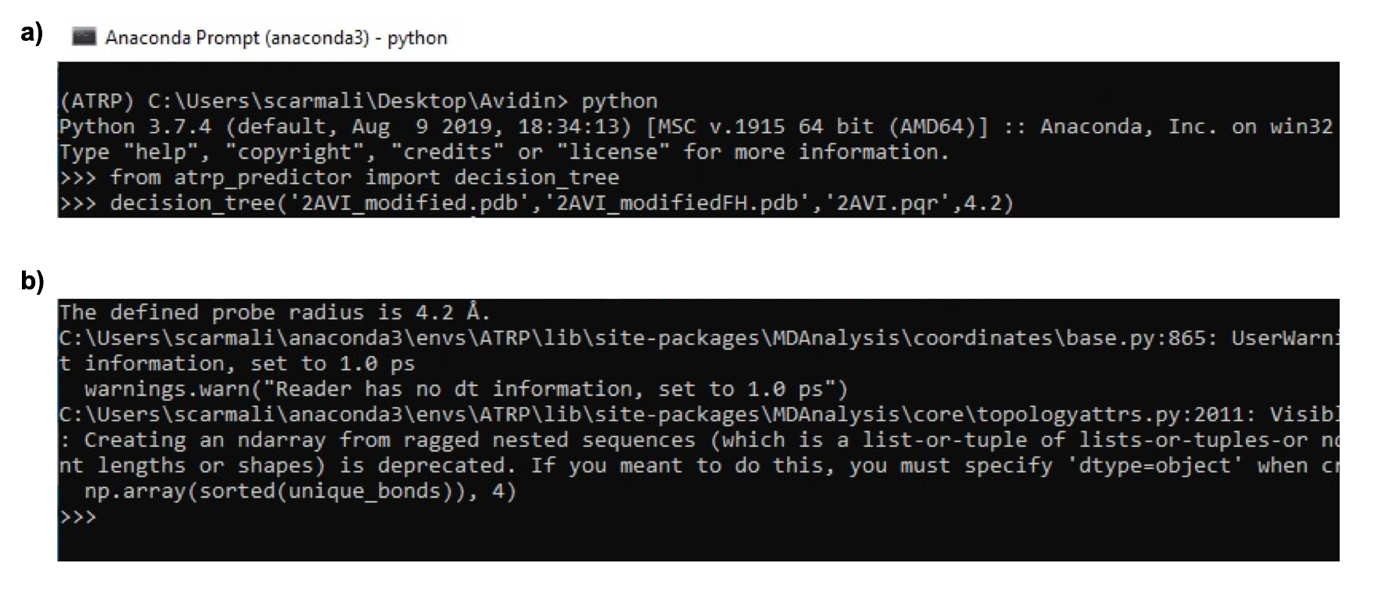

S3 Fig. PRELYM execution using Python 3.7.4 in the Anaconda Prompt. In the Anaconda Prompt, PRELYM is executed by a) calling the decision tree function with the required arguments. Upon running, PRELYM will b) confirm the defined probe radius and output a series of user warnings before generating a ‘results’ spreadsheet file in the common folder.

S1 Table PRELYM results for amine-ATRP initiator interactions on the surface of lysozyme. Shaded in grey are the experimental reactivity data for lysozyme from site modification studies with *N*-hydroxysuccinimide ATRP initiator.[2]

| **Chain** | **Residue** | **-NH2 Group** | **ESA (Å^2^)** | **pKa** | **Secondary Structure** | **H-Donor** | **Area of Lower Charge** | **Reactivity** | |
| --- | --- | --- | --- | --- | --- | --- | --- | --- | --- |
|  |  |  |  |  |  |  |  | **Predicted** | **Experimental** |
| A | K1 | α | 136.34 | 7.43 | Coil | No | Yes | fast-reacting | *not determined* |
|  | K1 | ε | 136.34 | 11.40 | Coil | No | Yes | fast-reacting | *not determined* |
|  | K13 | ε | 69.02 | 11.54 | Helix | Yes | Yes | slow-reacting | slow-reacting |
|  | K33 | ε | 75.91 | 10.14 | Helix | Yes | Yes | slow-reacting | slow-reacting |
|  | K96 | ε | 38.36 | 10.09 | Helix | Yes | Yes | non-reacting | non-reacting |
|  | K97 | ε | 145.03 | 10.45 | Helix | No | Yes | slow-reacting | slow-reacting |
|  | K116 | ε | 134.68 | 10.06 | Coil | Yes | Yes | fast-reacting | fast-reacting |

S2 Table PRELYM results for amine interactions on the surface of lysozyme using a probe radius equivalent to the hydrodynamic radius of *N*-hydroxysuccinimide RAFT CTA (7 Å; see S4 Fig.). Shaded in grey are experimental data for lysozyme from site modification studies with *N*-hydroxysuccinimide RAFT CTA.[3]

| **Chain** | **Residue** | **-NH2 Group** | **ESA (Å^2^)** | **pKa** | **Secondary Structure** | **H-Donor** | **Area of Lower Charge** | **Reactivity** | |
| --- | --- | --- | --- | --- | --- | --- | --- | --- | --- |
|  |  |  |  |  |  |  |  | **Predicted** | **Experimental** |
| A | K1 | α | 173.27 | 7.43 | Coil | No | Yes | fast-reacting | *not determined* |
|  | K1 | ε | 173.27 | 11.40 | Coil | No | Yes | fast-reacting | *not determined* |
|  | K13 | ε | 70.84 | 11.54 | Helix | Yes | Yes | slow-reacting | *not determined* |
|  | K33 | ε | 95.83 | 10.14 | Helix | Yes | Yes | slow-reacting | modified |
|  | K96 | ε | 35.64 | 10.09 | Helix | Yes | Yes | non-reacting | *not determined* |
|  | K97 | ε | 174.27 | 10.45 | Helix | No | Yes | slow-reacting | modified |
|  | K116 | ε | 185.04 | 10.06 | Coil | Yes | Yes | fast-reacting | *not determined* |

S3 Table PRELYM results for amine interactions on the surface of lysozyme using a probe radius equivalent to the hydrodynamic radius of oligomeric RAFT CTA (8.8 Å; see S4 Fig.). Shaded in grey are experimental data for lysozyme from site modification studies with oligomeric RAFT CTA.[4]

| **Chain** | **Residue** | **-NH2 Group** | **ESA (Å^2^)** | **pKa** | **Secondary Structure** | **H-Donor** | **Area of Lower Charge** | **Reactivity** | |
| --- | --- | --- | --- | --- | --- | --- | --- | --- | --- |
|  |  |  |  |  |  |  |  | **Predicted** | **Experimental** |
| A | K1 | α | 205.02 | 7.43 | Coil | No | Yes | fast-reacting | modified |
|  | K1 | ε | 205.02 | 11.40 | Coil | No | Yes | fast-reacting | *not determined* |
|  | K13 | ε | 64.43 | 11.54 | Helix | Yes | Yes | slow-reacting | *not determined* |
|  | K33 | ε | 112.11 | 10.14 | Helix | Yes | Yes | slow-reacting | modified |
|  | K96 | ε | 34.66 | 10.09 | Helix | Yes | Yes | non-reacting | *not determined* |
|  | K97 | ε | 195.73 | 10.45 | Helix | No | Yes | slow-reacting | modified |
|  | K116 | ε | 218.96 | 10.06 | Coil | Yes | Yes | fast-reacting | *not determined* |

S4 Table PRELYM results for amine interactions on the surface of lysozyme using a probe radius approximate to the hydrodynamic radius of PEG 5 kDa[5] (17 Å). Shaded in grey are experimental data for lysozyme from site modification studies with *N*-hydroxysuccinimide PEG reagent 5 kDa.[6]

| **Chain** | **Residue** | **-NH2 Group** | **ESA (Å^2^)** | **pKa** | **Secondary Structure** | **H-Donor** | **Area of Lower Charge** | **Reactivity** | |
| --- | --- | --- | --- | --- | --- | --- | --- | --- | --- |
|  |  |  |  |  |  |  |  | **Predicted** | **Experimental** |
| A | K1 | α | 294.85 | 7.43 | Coil | No | Yes | fast-reacting | modified |
|  | K1 | ε | 294.85 | 11.40 | Coil | No | Yes | fast-reacting | *modified** |
|  | K13 | ε | 64.87 | 11.54 | Helix | Yes | Yes | slow-reacting | *modified* |
|  | K33 | ε | 177.79 | 10.14 | Helix | Yes | Yes | slow-reacting | modified |
|  | K96 | ε | 26.33 | 10.09 | Helix | Yes | Yes | non-reacting | *not determined* |
|  | K97 | ε | 285.34 | 10.45 | Helix | No | Yes | slow-reacting | modified |
|  | K116 | ε | 361.61 | 10.06 | Coil | Yes | Yes | fast-reacting | *modified* |

* Only one amine site at K1 is modified.

S5 Table PRELYM results for amine-ATRP initiator interactions on the surface of homo-dimer chymotrypsin. Shaded in grey are the experimental reactivity data for chymotrypsin from site modification studies with *N*-hydroxysuccinimide ATRP initiator.[2]

| **Chain** | **Residue** | **-NH2 Group** | **ESA (Å^2^)** | **pKa** | **Secondary Structure** | **H-Donor** | **Area of Lower Charge** | **Reactivity** | |
| --- | --- | --- | --- | --- | --- | --- | --- | --- | --- |
|  |  |  |  |  |  |  |  | **Predicted** | **Experimental** |
| A | C1 | α | 126.0 | 7.66 |  | No |  | fast-reacting | fast-reacting |
| B | I16 | α | 0 |  |  | No |  | non-reacting | *not determined* |
|  | K36 | ε | 131.57 | 10.46 | Coil | No | No | slow-reacting* | fast-reacting |
|  | K79 | ε | 275.10 | 10.45 | Coil | No | Yes | fast-reacting | slow-reacting |
|  | K82 | ε | 120.04 | 10.36 | Strand | No | No | slow-reacting | non-reacting |
|  | K84 | ε | 184.79 | 10.41 | Strand | No | Yes | slow-reacting | *not determined* |
|  | K87 | ε | 181.62 | 10.24 | Strand | Yes | Yes | fast-reacting | *not determined* |
|  | K90 | ε | 86.98 | 10.12 | Strand | No | Yes | fast-reacting | slow-reacting |
|  | K93 | ε | 217.84 | 10.41 | Coil | Yes | Yes | fast-reacting | *not determined* |
|  | K107 | ε | 53.90 | 10.79 | Strand | Yes | No | slow-reacting | non-reacting |
| C | A149 | α | 5.50 | 7.54 |  | No |  | non-reacting | *not determined* |
|  | K169 | ε | 67.26 | 10.37 | Helix | Yes | Yes | slow-reacting | slow-reacting |
|  | K170 | ε | 286.60 | 10.49 | Helix | Yes | Yes | non-reacting | non-reacting |
|  | K175 | ε | 32.42 | 9.89 | Helix | Yes | Yes | non-reacting | *not determined* |
|  | K177 | ε | 57.55 | 10.13 | Coil | Yes | Yes | slow-reacting | slow-reacting |
|  | K202 | ε | 104.07 | 10.22 | Strand | No | Yes | fast-reacting | fast-reacting |
|  | K203 | ε | 58.88 | 10.74 | Strand | Yes | Yes | non-reacting | non-reacting |

S6 Table PRELYM results for amine-ATRP initiator interactions on the surface of monomer chymotrypsin. Shaded in grey are the experimental reactivity data for chymotrypsin from site modification studies with *N*-hydroxysuccinimide ATRP initiator.[2]

| **Chain** | **Residue** | **-NH2 Group** | **ESA (Å^2^)** | **pKa** | **Secondary Structure** | **H-Donor** | **Area of Lower Charge** | **Reactivity** | |
| --- | --- | --- | --- | --- | --- | --- | --- | --- | --- |
|  |  |  |  |  |  |  |  | **Predicted** | **Experimental** |
| A | C1 | α | 126.0 | 7.66 |  | No |  | fast-reacting | fast-reacting |
| B | I16 | α | 0 |  |  | No |  | non-reacting | *not determined* |
|  | K36 | ε | 221.63 | 10.45 | Coil | No | No | slow-reacting* | fast-reacting |
|  | K79 | ε | 275.10 | 10.45 | Coil | No | Yes | fast-reacting | slow-reacting |
|  | K82 | ε | 120.04 | 10.36 | Strand | No | No | slow-reacting | non-reacting |
|  | K84 | ε | 184.79 | 10.41 | Strand | No | Yes | slow-reacting | *not determined* |
|  | K87 | ε | 181.62 | 10.24 | Strand | Yes | Yes | fast-reacting | *not determined* |
|  | K90 | ε | 117.85 | 10.33 | Strand | No | Yes | slow-reacting | slow-reacting |
|  | K93 | ε | 217.84 | 10.41 | Coil | Yes | Yes | fast-reacting | *not determined* |
|  | K107 | ε | 53.90 | 10.79 | Strand | Yes | No | slow-reacting | non-reacting |
| C | A149 | α | 261.37 | 7.89 |  | No |  | fast-reacting | *not determined* |
|  | K169 | ε | 67.26 | 10.37 | Helix | Yes | Yes | slow-reacting | slow-reacting |
|  | K170 | ε | 290.49 | 10.49 | Helix | Yes | Yes | non-reacting | non-reacting |
|  | K175 | ε | 123.07 | 10.25 | Helix | Yes | Yes | slow-reacting | *not determined* |
|  | K177 | ε | 57.55 | 10.10 | Coil | Yes | Yes | slow-reacting | slow-reacting |
|  | K202 | ε | 104.07 | 10.22 | Strand | No | Yes | fast-reacting | fast-reacting |
|  | K203 | ε | 58.88 | 10.74 | Strand | Yes | Yes | non-reacting | non-reacting |

S7 Table PRELYM results for amine interactions on the surface of monomer chymotrypsin using a probe radius approximate to the hydrodynamic radius of PEG 5 kDa [5] (17 Å).

| **Chain** | **Residue** | **-NH2 Group** | **ESA (Å^2^)** | **pKa** | **Secondary Structure** | **H-Donor** | **Area of Lower Charge** |  |
| --- | --- | --- | --- | --- | --- | --- | --- | --- |
|  |  |  |  |  |  |  |  | **Predicted** |
| A | C1 | α | 169.61 | 7.66 |  | No |  | fast-reacting |
| B | I16 | α | 0 |  |  | No |  | non-reacting |
|  | K36 | ε | 673.34 | 10.45 | Coil | No | No | slow-reacting* |
|  | K79 | ε | 754.37 | 10.45 | Coil | No | Yes | fast-reacting |
|  | K82 | ε | 46.57 | 10.36 | Strand | No | No | non-reacting |
|  | K84 | ε | 439.01 | 10.41 | Strand | No | Yes | slow-reacting |
|  | K87 | ε | 421.02 | 10.24 | Strand | Yes | Yes | fast-reacting |
|  | K90 | ε | 131.41 | 10.33 | Strand | No | Yes | slow-reacting |
|  | K93 | ε | 538.57 | 10.41 | Coil | Yes | Yes | fast-reacting |
|  | K107 | ε | 39.50 | 10.79 | Strand | Yes | No | non-reacting |
| C | A149 | α | 627.86 | 7.89 |  | No |  | fast-reacting |
|  | K169 | ε | 17.52 | 10.37 | Helix | Yes | Yes | non-reacting |
|  | K170 | ε | 1046.5 | 10.49 | Helix | Yes | Yes | slow-reacting |
|  | K175 | ε | 179.85 | 10.25 | Helix | Yes | Yes | slow-reacting |
|  | K177 | ε | 9.38 | 10.10 | Coil | Yes | Yes | non-reacting |
|  | K202 | ε | 86.31 | 10.22 | Strand | No | Yes | fast-reacting |
|  | K203 | ε | 4.52 | 10.74 | Strand | Yes | Yes | non-reacting |

S8 Table PRELYM results for amine-ATRP initiator interactions on the surface of monomer glucose oxidase.

| **Chain** | **Residue** | **-NH2 Group** | **ESA (Å^2^)** | **pKa** | **Secondary Structure** | **H-Donor** | **Area of Lower Charge** | **Predicted**  **Reactivity** |
| --- | --- | --- | --- | --- | --- | --- | --- | --- |
| A | S1 | α | 320.65 | 7.90 |  | No |  | fast-reacting |
|  | K13 | ε | 210.81 | 10.35 | Helix | Yes | No | slow-reacting |
|  | K116 | ε | 97.34 | 10.40 | Helix | Yes | No | slow-reacting |
|  | K152 | ε | 50.32 | 11.31 | Helix | Yes | No | slow-reacting |
|  | K187 | ε | 97.76 | 10.19 | Helix | No | No | slow-reacting |
|  | K201 | ε | 56.25 | 9.79 | Coil | Yes | No | slow-reacting |
|  | K202 | ε | 98.19 | 10.42 | Coil | No | No | non-reacting |
|  | K252 | ε | 38.57 | 10.22 | Strand | No | No | non-reacting |
|  | K273 | ε | 196.68 | 10.54 | Coil | No | No | slow-reacting |
|  | K282 | ε | 142.90 | 10.58 | Strand | Yes | No | slow-reacting |
|  | K306 | ε | 176.18 | 10.44 | Helix | Yes | No | slow-reacting |
|  | K364 | ε | 56.75 | 10.08 | Helix | No | No | slow-reacting |
|  | K372 | ε | 68.69 | 10.48 | Helix | Yes | No | slow-reacting |
|  | K441 | ε | 137.09 | 10.30 | Coil | Yes | No | fast-reacting |
|  | K526 | ε | 90.31 | 10.13 | Helix | No | No | slow-reacting |
|  | K570 | ε | 0 | 8.61 | Helix | Yes | No | non-reacting |

S9 Table PRELYM results for amine-ATRP initiator interactions on the surface of dimer avidin. Shaded in grey are the experimental data for avidin from site modification studies with *N*-hydroxysuccinimide ATRP initiator.[7]

| **Chain** | **Residue** | **-NH2 Group** | **ESA (Å^2^)** | **pKa** | **Secondary Structure** | **H-Donor** | **Area of Lower Charge** | **Predicted** | **Experimental Modification** |
| --- | --- | --- | --- | --- | --- | --- | --- | --- | --- |
|  |  |  |  |  |  |  |  | **Reactivity** |  |
| A | A1 | α | 84.86 | 6.65 |  | No |  | fast-reacting | *not determined* |
|  | K3 | ε | 229.99 | 10.41 | Coil | No | Yes | fast-reacting | *not determined* |
|  | K9 | ε | 41.99 | 10.28 | Strand | No | Yes | non-reacting | *not determined* |
|  | K45 | ε | 143.07 | 10.39 | Coil | No | Yes | fast-reacting | modified |
|  | K58 | ε | 210.76 | 10.20 | Coil | No | Yes | fast-reacting | *not determined* |
|  | K71 | ε | 137.21 | 10.12 | Coil | Yes | Yes | fast-reacting | modified |
|  | K90 | ε | 123.78 | 10.47 | Coil | No | Yes | fast-reacting | *not determined* |
|  | K94 | ε | 18.43 | 10.02 | Strand | Yes | Yes | non-reacting | *not determined* |
|  | K111 | ε | 120.41 | 10.39 | Helix | No | Yes | slow-reacting | modified |
|  | K127 | ε | 286.97 | 10.46 | Coil | No |  | slow-reacting | *not determined* |
| B | A1 | α | 114.08 | 7.68 |  | No |  | fast-reacting | *not determined* |
|  | K3 | ε | 136.65 | 10.45 | Coil | No | Yes | fast-reacting | *not determined* |
|  | K9 | ε | 172.78 | 10.35 | Strand | No | Yes | slow-reacting | *not determined* |
|  | K45 | ε | 151.79 | 10.55 | Coil | No | Yes | fast-reacting | modified |
|  | K58 | ε | 152.60 | 10.11 | Coil | No | Yes | fast-reacting | *not determined* |
|  | K71 | ε | 187.98 | 10.26 | Coil | Yes | Yes | fast-reacting | modified |
|  | K90 | ε | 70.92 | 10.36 | Coil | No | Yes | slow-reacting | *not determined* |
|  | K94 | ε | 30.96 | 10.23 | Strand | Yes | Yes | non-reacting | *not determined* |
|  | K111 | ε | 118.67 | 10.27 | Helix | No | Yes | slow-reacting | modified |
|  | K127 | ε | 329.13 | 10.47 | Coil | No |  | slow-reacting | *not determined* |

S10 Table PRELYM results for amine-ATRP initiator interactions on the surface of interferon-α 2a. Shaded in grey are experimental data obtained from site modification studies with PEGylated interferon-α 2a using an amine-reactive branched 40 kDa PEG.[8]

| **Chain** | **Residue** | **-NH2 Group** | **ESA (Å^2^)** | **pKa** | **Secondary Structure** | **H-Donor** | **Area of Lower Charge** | **Predicted**  **Reactivity** | **PEGylation Sites in INF-α 2a** |
| --- | --- | --- | --- | --- | --- | --- | --- | --- | --- |
| A | C1 | α | 36.01 | 7.68 |  | No |  | non-reacting | not modified |
|  | K23 | ε | 103.84 | 10.67 | Coil | Yes | No | fast-reacting | not modified |
|  | K31 | ε | 287.07 | 10.44 | Coil | Yes | Yes | fast-reacting | modified |
|  | K49 | ε | 187.51 | 11.16 | Coil | Yes | No | fast-reacting | modified |
|  | K70 | ε | 276.63 | 10.62 | Helix | No | Yes | slow-reacting | modified |
|  | K83 | ε | 94.70 | 10.29 | Helix | Yes | Yes | slow-reacting | modified |
|  | K112 | ε | 7.32 | 10.09 | Helix | No | No | non-reacting | modified |
|  | K121 | ε | 134.93 | 10.36 | Helix | No | No | slow-reacting | modified |
|  | K131 | ε | 143.79 | 11.56 | Helix | No | Yes | slow-reacting | modified |
|  | K133 | ε | 29.34 | 11.09 | Coil | Yes | Yes | non-reacting | not modified |
|  | K134 | ε | 304.32 | 10.42 | Coil | Yes | Yes | fast-reacting | modified |
|  | K164 | ε | 172.67 | 10.37 | Coil | No | Yes | fast-reacting | modified |

S11 Table PRELYM results for amine interactions on the surface of interferon-a 2a using a probe radius equivalent to the hydrodynamic radius of PEG 40 kDa (39.5 Å). Shaded in grey are experimental data obtained from site modification studies with PEGylated interferon-a 2a using an amine-reactive branched 40 kDa PEG.[8]

| **Chain** | **Residue** | **-NH2 Group** | **ESA (Å^2^)** | **pKa** | **Secondary Structure** | **H-Donor** | **Area of Lower Charge** | **Predicted**  **Reactivity** | **PEGylation Sites in INF-α 2a** |
| --- | --- | --- | --- | --- | --- | --- | --- | --- | --- |
| A | C1 | α | 0 | 7.68 |  | No |  | non-reacting | not modified |
|  | K23 | ε | 262.59 | 10.67 | Coil | Yes | No | fast-reacting | not modified |
|  | K31 | ε | 3153.57 | 10.44 | Coil | Yes | Yes | fast-reacting | modified |
|  | K49 | ε | 607.46 | 11.16 | Coil | Yes | No | fast-reacting | modified |
|  | K70 | ε | 2779.62 | 10.62 | Helix | No | Yes | slow-reacting | modified |
|  | K83 | ε | 90.00 | 10.29 | Helix | Yes | Yes | slow-reacting | modified |
|  | K112 | ε | 0 | 10.09 | Helix | No | No | non-reacting | modified |
|  | K121 | ε | 306.74 | 10.36 | Helix | No | No | slow-reacting | modified |
|  | K131 | ε | 252.83 | 11.56 | Helix | No | Yes | slow-reacting | modified |
|  | K133 | ε | 0 | 11.09 | Coil | Yes | Yes | non-reacting | not modified |
|  | K134 | ε | 2881.76 | 10.42 | Coil | Yes | Yes | fast-reacting | modified |
|  | K164 | ε | 0 | 10.37 | Coil | No | Yes | non-reacting | modified |

S12 Table PRELYM results for amine-ATRP initiator interactions on the surface of asparaginase II.

| **Chain** | **Residue** | **-NH2 Group** | **ESA (Å^2^)** | **pKa** | **Secondary Structure** | **H-Donor** | **Area of Lower Charge** | **Predicted**  **Reactivity** |
| --- | --- | --- | --- | --- | --- | --- | --- | --- |
| A | L1 | α | 207.88 | 7.85 |  | No |  | fast-reacting |
|  | K22 | ε | 122.08 | 10.47 | Coil | No | No | slow-reacting |
|  | K29 | ε | 137.77 | 10.34 | Coil | Yes | No | fast-reacting |
|  | K43 | ε | 128.70 | 10.28 | Helix | No | No | slow-reacting |
|  | K49 | ε | 92.84 | 10.38 | Strand | Yes | No | slow-reacting |
|  | K71 | ε | 22.037 | 9.99 | Helix | Yes | No | non-reacting |
|  | K72 | ε | 30.52 | 12.22 | Helix | Yes | No | non-reacting |
|  | K79 | ε | 155.17 | 10.97 | Helix | Yes | No | slow-reacting |
|  | K104 | ε | 46.36 | 8.88 | Coil | Yes | No | non-reacting |
|  | K107 | ε | 56.99 | 10.26 | Coil | Yes | No | slow-reacting |
|  | K139 | ε | 268.35 | 10.47 | Helix | No | No | slow-reacting |
|  | K162 | ε | 0 | 10.11 | Strand | Yes | No | non-reacting |
|  | K172 | ε | 1.26 | 10.36 | Strand | Yes | No | non-reacting |
|  | K186 | ε | 91.10 | 10.74 | Strand | Yes | No | slow-reacting |
|  | K196 | ε | 124.97 | 11.42 | Coil | Yes | No | fast-reacting |
|  | K207 | ε | 315.16 | 10.50 | Coil | No | No | slow-reacting |
|  | K213 | ε | 71.43 | 10.63 | Coil | Yes | No | slow-reacting |
|  | K229 | ε | 93.55 | 10.81 | Helix | Yes | No | slow-reacting |
|  | K251 | ε | 133.65 | 10.38 | Helix | No | No | slow-reacting |
|  | K262 | ε | 194.76 | 10.26 | Helix | No | No | slow-reacting |
|  | K288 | ε | 197.36 | 11.17 | Helix | No | No | slow-reacting |
|  | K301 | ε | 0 | 8.28 | Helix | Yes | No | non-reacting |
|  | K314 | ε | 166.0 | 10.51 | Coil | No | No | slow-reacting |

S13 Table PRELYM results for amine interactions on the surface of asparaginase II using a probe radius approximate to the hydrodynamic radius of PEG 5 kDa [9] (17 Å).

| **Chain** | **Residue** | **-NH2 Group** | **ESA (Å^2^)** | **pKa** | **Secondary Structure** | **H-Donor** | **Area of Lower Charge** | **Predicted**  **Reactivity** |
| --- | --- | --- | --- | --- | --- | --- | --- | --- |
| A | L1 | α | 419.19 | 7.85 |  | No |  | fast-reacting |
|  | K22 | ε | 75.66 | 10.47 | Coil | No | No | slow-reacting |
|  | K29 | ε | 132.54 | 10.34 | Coil | Yes | No | fast-reacting |
|  | K43 | ε | 233.36 | 10.28 | Helix | No | No | slow-reacting |
|  | K49 | ε | 67.31 | 10.38 | Strand | Yes | No | slow-reacting |
|  | K71 | ε | 0 | 9.99 | Helix | Yes | No | non-reacting |
|  | K72 | ε | 1.88 | 12.22 | Helix | Yes | No | non-reacting |
|  | K79 | ε | 434.17 | 10.97 | Helix | Yes | No | slow-reacting |
|  | K104 | ε | 0 | 8.88 | Coil | Yes | No | non-reacting |
|  | K107 | ε | 5.57 | 10.26 | Coil | Yes | No | non-reacting |
|  | K139 | ε | 509.46 | 10.47 | Helix | No | No | slow-reacting |
|  | K162 | ε | 0 | 10.11 | Strand | Yes | No | non-reacting |
|  | K172 | ε | 0 | 10.36 | Strand | Yes | No | non-reacting |
|  | K186 | ε | 0 | 10.74 | Strand | Yes | No | non-reacting |
|  | K196 | ε | 136.21 | 11.42 | Coil | Yes | No | fast-reacting |
|  | K207 | ε | 972.58 | 10.50 | Coil | No | No | slow-reacting |
|  | K213 | ε | 0 | 10.63 | Coil | Yes | **No** | non-reacting |
|  | K229 | ε | 195.24 | 10.81 | Helix | Yes | No | slow-reacting |
|  | K251 | ε | 167.62 | 10.38 | Helix | No | No | slow-reacting |
|  | K262 | ε | 351.93 | 10.26 | Helix | No | No | slow-reacting |
|  | K288 | ε | 419.62 | 11.17 | Helix | No | No | slow-reacting |
|  | K301 | ε | 0 | 8.28 | Helix | Yes | No | non-reacting |
|  | K314 | ε | 349.18 | 10.51 | Coil | No | No | slow-reacting |

S14 Table PRELYM results for amine-ATRP initiator interactions on the surface of tetrameric phenylalanine ammonia lyase (rAV-PAL). Shaded in grey are experimental data obtained for rAV-PAL from site modification studies using a 20 kDa *N*-hydroxysuccinimide PEG.[10] Percentage of modification was determined by mass spectrometry after tryptic digestion.

| **Chain** | **Residue** | **-NH2 Group** | **ESA (Å^2^)** | **pKa** | **Secondary Structure** | **H-Donor** | **Area of Lower Charge** | **Predicted**  **Reactivity** | **PEGylation Sites**  **(% of modification)** |
| --- | --- | --- | --- | --- | --- | --- | --- | --- | --- |
|  | M1 | α | 468.51 | 7.93 |  | No |  | fast-reacting | *not determined* |
|  | K2 | ε | 270.99 | 10.41 | Coil | No |  | non-reacting | modified (100%) |
|  | K10 | ε | 282.00 | 10.46 | Coil | No |  | slow-reacting | modified (100%) |
|  | K32 | ε | 111.01 | 9.93 | Coil | No | No | slow-reacting | modified (40%) |
|  | K109 | ε | 0 | 10.18 | Coil | Yes | No | non-reacting | *not determined* |
|  | K115 | ε | 46.99 | 7.95 | Strand | No | No | non-reacting | modified (20%) |
|  | K145 | ε | 65.75 | 10.52 | Helix | Yes | No | slow-reacting | modified (50%) |
|  | K189 | ε | 38.13 | 10.44 | Strand | Yes | No | non-reacting | *not determined* |
|  | K195 | ε | 254.07 | 10.41 | Strand | Yes | No | slow-reacting | modified (100%) |
|  | K216 | ε | 0.109 | 9.18 | Coil | Yes | No | non-reacting | *not determined* |
|  | K272 | ε | 0 | 9.55 | Coil | Yes | No | non-reacting | *not determined* |
|  | K301 | ε | 171.12 | 10.60 | Coil | Yes | No | fast-reacting | modified (40%) |
|  | K335 | ε | 107.23 | 10.10 | Helix | Yes | No | slow-reacting | modified (20%) |
|  | K384 | ε | 0 | 5.89 | Helix | Yes | No | non-reacting | *not determined* |
|  | K413 | ε | 170.57 | 10.37 | Coil | No | No | slow-reacting | modified (90%) |
|  | K419 | ε | 0 | 10.68 | Helix | Yes | No | non-reacting | modified (20%) |
|  | K493 | ε | 147.60 | 9.98 | Helix | Yes | No | slow-reacting | *modified (100%) |
|  | K494 | ε | 124.47 | 10.34 | Helix | Yes | No | non-reacting | *modified (100%) |
|  | K522 | ε | 230.93 | 10.42 | Coil | No | No | slow-reacting | modified (100%) |

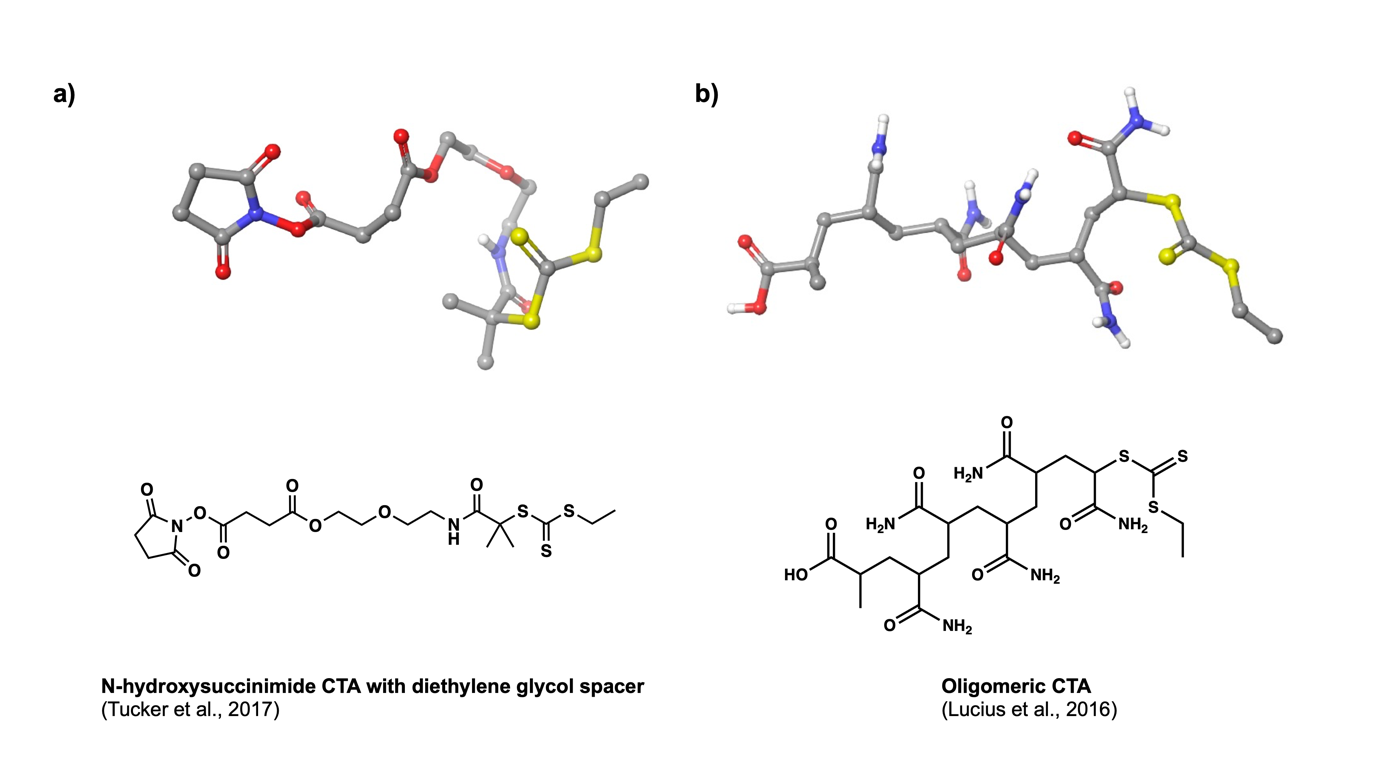

S4 Fig. Chain transfer agents used for surface-initiated RAFT of lysozyme. 3D and chemical structures of a) *N*-hydroxysuccinimide CTA with diethylene glycol spacer used for photoinduced electron/energy transfer-reversible addition-fragmentation chain transfer (PET-RAFT) polymerisation and b) oligomeric RAFT CTA with 5 acrylamide units.

**Estimation of probe radius for chain transfer agents.** CTA chemical structures were first built using Maestro 2D Sketcher and imported into the main workspace area for 3D transformation.[11] Geometry minimisation was carried out using the default force field (OPLS_2005).[12] End-to-end distances were determined using the “measure” tool in Maestro. The obtained distance was considered as the theoretical diameter of the corresponding CTA, and the value divided by 2 was used as the probe radius for PRELYM calculations.
